# Spatially Resolved and Highly Complexed Protein and RNA *in situ* Detection by Combining CODEX with RNAscope In Situ Hybridization

**DOI:** 10.1101/2022.02.10.479971

**Authors:** Yilun Cheng, Rachel M. Burrack, Qingsheng Li

## Abstract

Highly multiplexed protein and RNA *in situ* concurrent detection on a single tissue section is highly desirable for both basic and applied biomedical research. CODEX is a new and powerful platform to visualize up to 60 protein biomarkers *in situ* and RNAscope *in situ* hybridization (RNAscope) is a novel RNA detection system with single-copy sensitivity and unprecedent specificity at a single cell level. Nevertheless, to our knowledge, the combination CODEX and RNAscope remained unreported until this study. Here we report a simple and reproducible combination of CODEX and RNAscope (Comb-CODEX-RNAscope). We also determined the cross-reactivities of CODEX anti-human antibodies to rhesus macaques, a widely used animal model of human disease.

## INTRODUCTION

Highly multiplexed protein *in situ* detection (HMPISD) on a single tissue section at subcellular resolution is a powerful new technology to visualize and quantify proteins in their native tissue microenvironment with spatial context (1–3). The HMPISD technologies have been wildly used in basic and applied biological research. The HMPISD has also been used in cancer biology to explore the complex immune landscapes and tumor microenvironment (1, 3–7) and in the study of pathogenesis of autoimmune disease (8). Several strategies and platforms of HMPISD have been developed and utilized including 1) Time-of-Flight Mass Spectrometry based platforms of Imagine Mass Cytometry (IMC) (9, 10) and Multiplexed Ion Beam Imaging (MIBI) (11, 12). Both methods combined metal-labeled antibody immunostaining, UV laser ablation in IMC or ion beam gun ablation in MIBI, and CyTOF mass cytometry; 2) iterative fluorescent-conjugated antibody staining, imaging, and removing antibody or inactivating fluorophores. For example, MultiOmyx (13–15) and the tissue-based CyClic ImmunoFluorescence (t-CyCIF) (16, 17); and 3) DNA-barcoded antibody platforms. For example, the Immunostaining with Signal Amplification by Exchange Reaction (Immuno-SABER) (18, 19) and CO-Detection by inDEXing (CODEX). In Immuno-SABER, tissue target proteins are first bound with a panel of DNA-conjugated antibodies, subsequently multiple rounds of hybridizing with corresponding concatemer DNA and fluorescent-DNA, imaging, and removing the fluorophores. CODEX is a new and powerful platform, of which multiple epitopes are first bound with a panel of barcoded oligonucleotide conjugated antibodies followed by multiple cycles of adding corresponding fluorescently-labeled-oligonucleotide probes, imaging, and removing fluorescently-labeled-oligonucleotide probes to visualize up to 60 protein biomarkers *in situ* (6, 8, 20). While the prototype CODEX using dNTP analogs plus DNA polymerase primer extension to amplify signal of DNA barcode conjugated to antibody (8), the current CODEX uses chaotropic solvents to facilitate sequential room temperature (RT) annealing and stripping process without polymerase reaction (6). Compared to other HMPISD methods, CODEX enlarges multiplexing ability (3, 20), increases detection resolution to subcellular level(1), preserves tissue architecture, and reduces experiment duration through single staining procedure(1). Simultaneous visualization of spatially resolved and highly multiplexed proteins and RNA *in situ* is highly desirable for both basic and applied biomedical research (21, 22). RNAscope *in situ* hybridization (RNAscope) from Advanced Cell Diagnostics (ACD) is a novel RNA detection system with single-copy sensitivity and unprecedent specificity at a single cell level (19, 23). Nevertheless, RNAscope and CODEX involve complex chemical and physical procedures. To our knowledge, the combination CODEX and RNAscope (Comb-CODEX-RNAscope) remained unreported until this study.

In this study, we tested various methods of combination and found a simple and reproducible approach of CODEX and subsequent RNAscope (Comb-CODEX-RNAscope) to visualize highly complexed proteins and RNA simultaneously in a single tissue section. Currently, highly multiplexed protein *in situ* detection platforms including CODEX is designed for human sample detection using anti-human antibodies. Non-human primates (NHP) evolutionally, anatomically, and physiologically are the closest species to humans and are regarded as one the best animal models of many human diseases. Nevertheless, the CODEX antibody compatibility to NHP was unknow until this study. We determined the cross-reactivities of CODEX anti-human antibodies to rhesus macaques, which are the most widely used NHP to model human disease including HIV (24, 25).

## RESULTS

### Determine the cross-reactivities of CODEX anti-human antibodies to rhesus macaques

To determine the cross-reactivities of CODEX anti-human antibodies to rhesus macaques, a panel of DNA-barcoded anti-human antibodies from Akoya Biosciences (Akoya) was tested in various fixative fixed (SafeFixII, 4% Paraformaldehyde, neutral-buffered formalin) and paraffin embedded lymph node tissues of rhesus macaques that were infected with Simian Immunodeficiency virus (SIV). We found 13 out of 24 CODEX anti-human antibodies from Akoya were reactive specifically with rhesus macaques (**Table 1 & Fig 1**). The DNA-barcoded anti-human CD3 antibody from Akoya (CD3e, Clone# EP449E) did not react with rhesus macaque tissues. We therefore conjugated an anti-human CD3 antibody (SP162, Cat#: ab245731, Abcam) with a DNA-barcoded oligonucleotide (Cat#: 5350002, Akoya) by following the conjugating kit manual (Cat#: 7000009, Akoya). After verifying the correct size through protein gel electrophoresis (data not shown), the conjugated CD3 antibody was tested using CODEX. It worked well for rhesus macaque tissues (**Fig 2**), as expected where CD3 signals were clearly separated from CD20 signals, partially colocalized with CD4 signals, and primarily distributed in T cell zone of lymph node tissues.

**Table 1.**
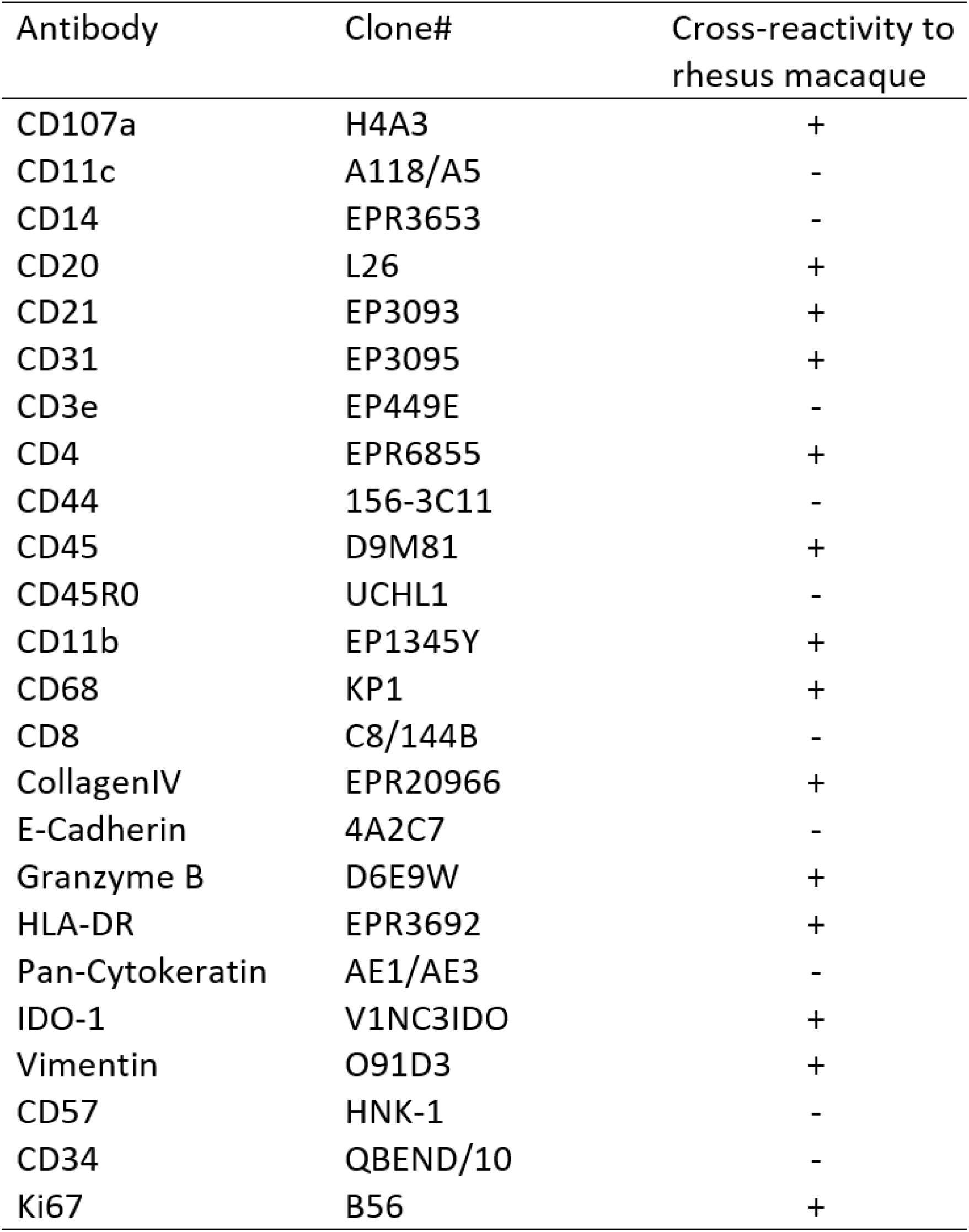
The cross-reactivities of CODEX anti-human DNA-Barcoded-antibodies from Akoya to rhesus macaques

**Figure 1.**
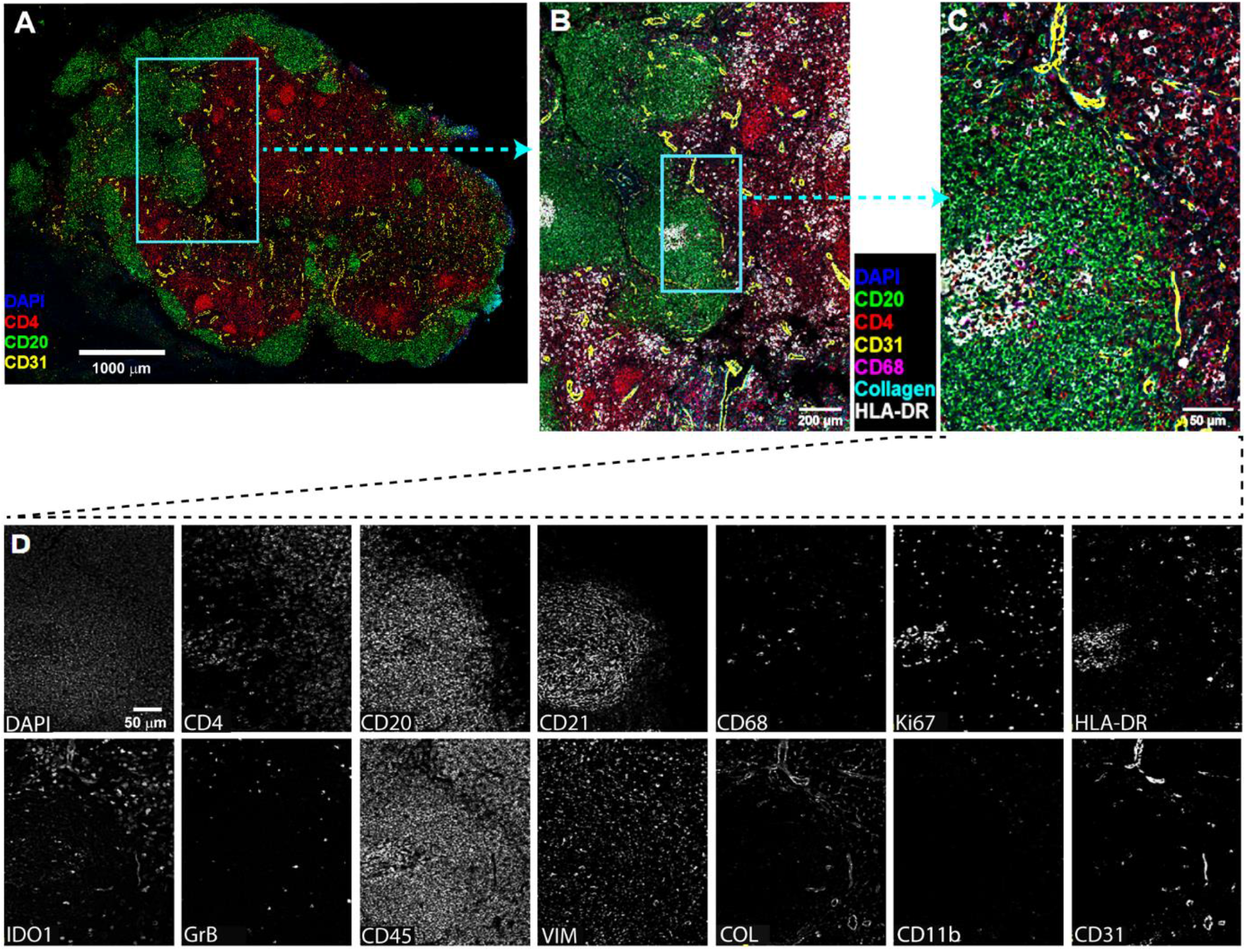
The cross-reactivities of 24 DNA-barcoded anti-human antibodies from Akoya to rhesus macaques using CODEX. (**A**) Overview image of lymph node tissue from a rhesus macaque infected with SIVmac251 (Rh4979, 10 dpi), showing CD4 (red), CD20 (green), DAPI (blue) and CD31(yellow). (**B**) The boxed area in A was zoomed to show DAPI (blue), CD4 (red), CD20 (green), CD68 (magenta), Collagen (cyan), HLA-DR (white) and CD31(yellow). (C) The boxed area in B was zoomed. (**D**) individual channel image showing 13 cross-reactive anti-human antibodies to rhesus macaque. Scale bars are shown.

**Figure 2.**
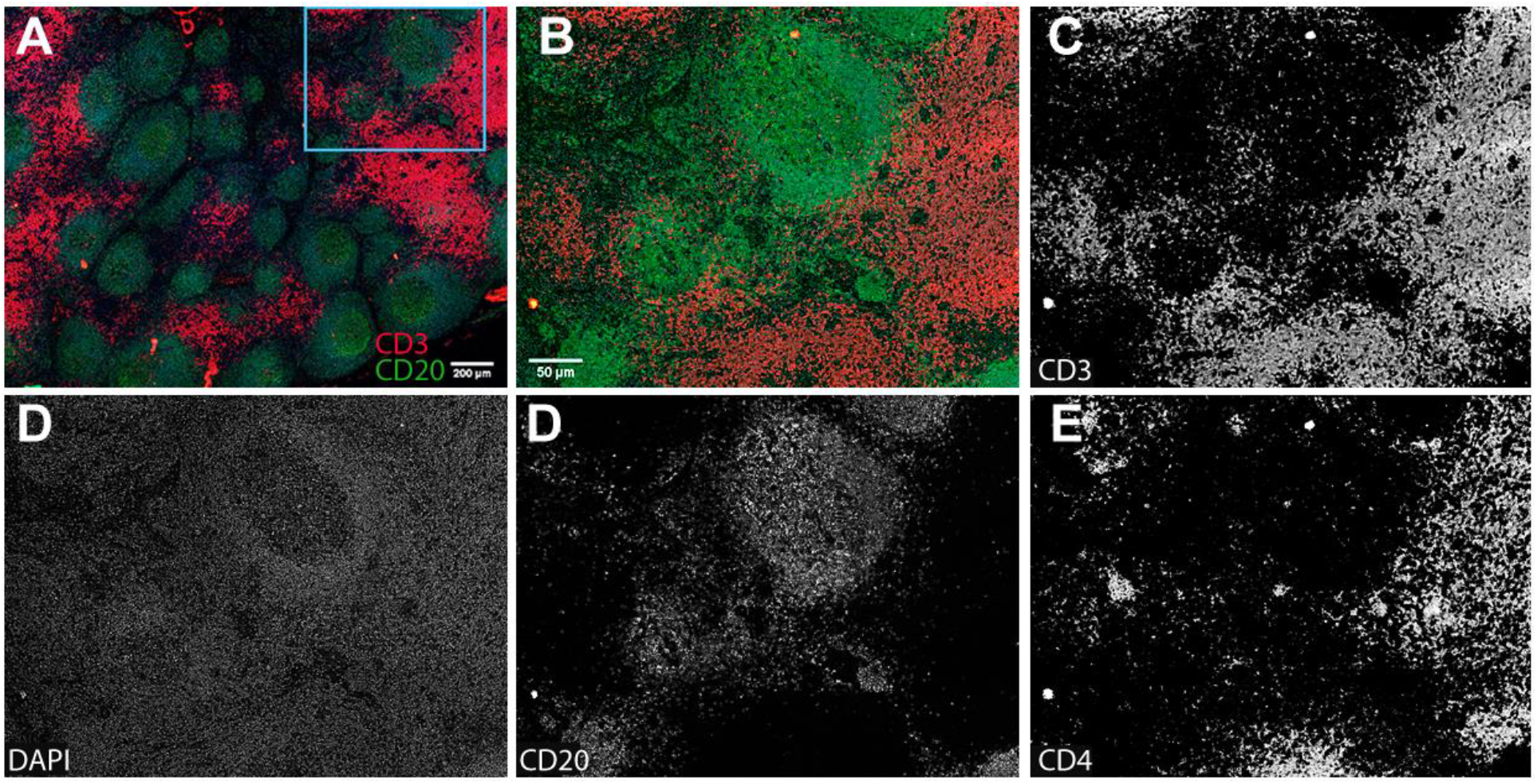
Validation of a home anti-human CD3 antibody (SP162, Cat#: ab245731, Abcam) conjugated with a DNA-barcode using CODEX to rhesus macaques. (**A**) Overview image of lymph node tissue from a rhesus macaque infected with SIVmac251 (Rh4979, 10 dpi). (**B**) The boxed area in A was zoomed. (**C-E**) individual channel image of antibodies, showing CD3, CD20, CD4 and DAPI. Scale bars are shown.

### The Combination of CODEX immunostaining with RNAscope ISH

After identifying rhesus macaque compatible antibodies, we selected a subset of them that are important for SIV immunopathogenesis to test various methods of combining CODEX with RNAscope. Two different orders of combination, RNAscope and subsequent CODEX as well as CODEX and subsequent RNAscope, were tested. All the procedures including deparaffinization, tissue hydration and antigen retrieval were conducted to minimize RNase contamination. RNAscope and subsequent CODEX method significantly induced non-specific signals and reduced the specific signals of CODEX. This combination led to unaccepted low signal-noise-ratio (SNR) of CODEX as compared with a single CODEX, although RNAscope signals maintained before and after CODEX (data not shown). Since RNAscope and subsequent CODEX approach could not get satisfying results, next efforts were focused on the approach of CODEX and subsequent RNAscope. When following the protocol of single CODEX and single RNAscope sequentially, RNAscope signals could not consistently detected, although CODEX signal detection worked well. It was plausible that some steps of CODEX including the antigen-retrieval step might impair RNAscope signal detection. It was tested and excluded the impact of number of cycles of CODEX on the RNAscope signal detection by comparing single cycle versus multiple cycles of CODEX immunostaining. It was further excluded the impact of RNase contamination of CODEX fluidics instrument by adding CODEX fluidics washing step into a single RNAscope procedure. The washing with CODEX fluidics did not impact RNAscope signal detection. It was narrowed down that the source of inconsistent RNAscope signal detection was in the antigen-retrieval step using a high-pressure cooker (Decloaking Chamber™, Biocare Medical). An alternative method for antigen retrieval using 50 ml glass-beaker on a Hotplate (Thermo Fisher, SP88857104) was tested, where the coverslips with tissue sections were placed in the coverslip staining rack (PN# 72240, Electron Microscopy Science) and immersed in boiling citrate buffer (pH 6, Sigma 21545) for 20 min. This method is simple and easy to eliminate potential RNase contamination by pre-baking glass-beaker and coverslip staining rack covered with aluminum foil at 300 ^o^C for 6 hrs. The developed CODEX and subsequent RNAscope approach (Comb-CODEX-RNAscope) with hotplate antigen retrieval is simple and reproducible for both highly multiplexed protein and RNA detections for SafeFixII, 4% paraformaldehyde, and neutral-buffered formalin fixed and paraffin-embedded tissues (**Fig 3 & 4**). SIV viral RNA (vRNA) signals from RNAscope (**Fig 4B-C**) in addition to CD3, CD4 CD68, CD20, CD21, CD31, HLA-DR, Ki67 and DAPI from CODEX (**Fig 4D-L**) were specific and intense enabling downstream spatial analyses.

**Figure 3.**
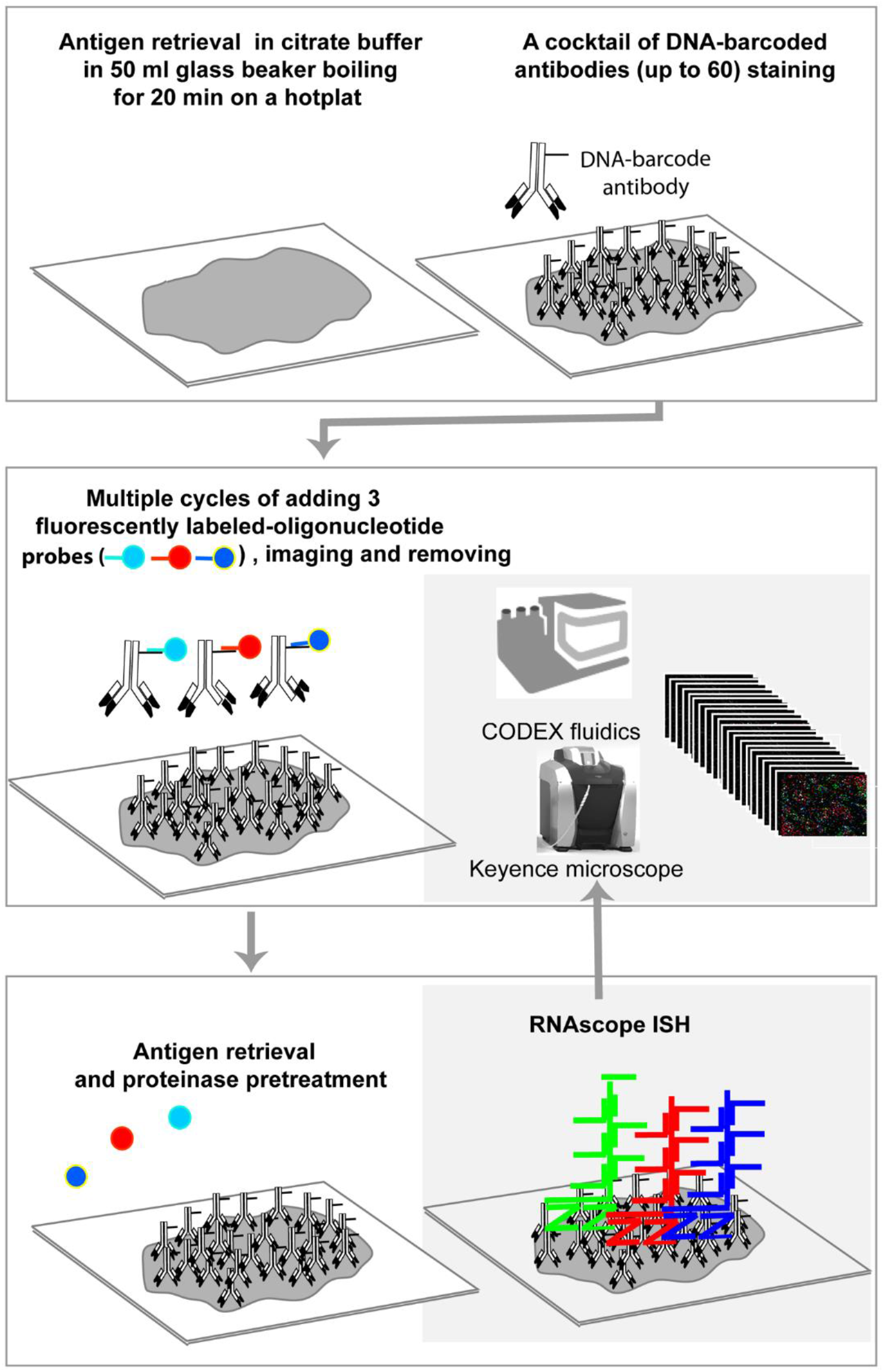
The workflow of CODEX and subsequent RNAscope in situ hybridization combination (comb-CODEX-RNAscope).

**Figure 4.**
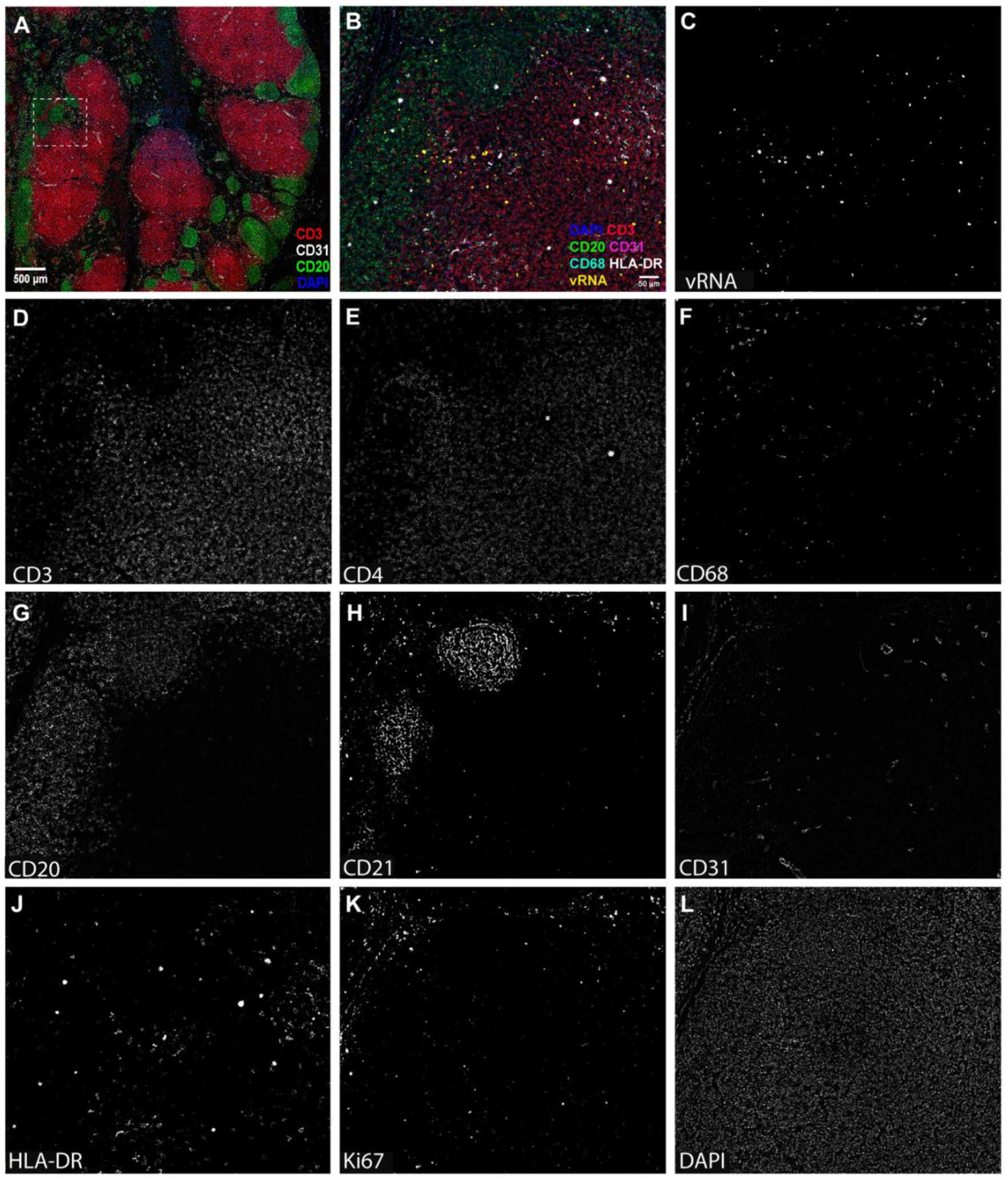
Representative images of the comb-CODEX-RNAscope. (**A**) Overview image of lymph node tissues from a rhesus macaque infected with SIVmac251 (Rh4979, 10 dpi), showing CD3 (red), CD20 (green), CD31(white) and DAPI (blue). (**B**) The boxed area in A was zoomed to show SIV viral RNA detected by RNAscope and other 5 protein markers by CODEX. (**C-L**) individual channel image showing vRNA and protein biomarkers. Scale bars are shown.

## MATERIALS AND METHODS

### Rhesus Macaque Tissues

Archived fixed lymphatic tissues from adult rhesus macaques infected with SIVmac251 from our previous reported studies were used(26, 27). Rhesus macaques were intra-rectally inoculated with SIVmac251 (3.1 x 10^4^ TCID_50_) and were euthanized at different days post-inoculation and rhesus macaques without virus inoculation served as uninfected controls. Lymph node tissues were fixed with SafeFix II (Cat# 23-042600, Fisher Scientific) for 6 hrs at RM, 4% paraformaldehyde 6 hrs at RT or neutral-buffered formalin for 24 hrs and embedded with paraffin.

### Antibody-oligonucleotide conjugation

To find a rhesus macaque reactive CD3 antibody for CODEX, an anti-human CD3 antibody in a carrier-free PBS solution (Clone#: SP162, Cat#: ab245731, Abcam) was conjugated with a barcode-oligonucleotide (Cat#: 5350002, Akoya) using CODEX conjugation kit (Cat#: 7000009, Akoya) following CODEX conjugation manual.

### CODEX immunostaining

CODEX immunostaining was performed according to CODEX user manual and previously published method (20). Six-μm tissue sections were cut using Leica RM2235 rotary microtome and mounted to poly-lysine coated coverslips. The following steps were performed in 6-well tissue culture plate. Tissue-coverslips were deparaffinized by sequentially heating on a hotplate at 60°C for I hr and immersed in xylene for 5 min twice and hydrated with gradually decreased concentration of ethanol and DEPC water. The antigen retrieval was performed in a high-pressure cooker (Decloaking Chamber™, Biocare Medical) for 20 min at high-pressure or Hotplate (Thermo Fisher, SP88857104) boiling for 20 min. The tissue-coverslips in the coverslip staining rack (PN# 72240, Electron Microscopy Science) were immersed in citrate buffer (pH 6, Sigma 21545) in 50 ml glass beaker. After antigen retrieval, the tissue-coverslips were stained with a cocktail of DNA-barcoded antibodies for 3 hrs at RT. After washing in PBS 3 times 2 min each to remove unbound antibodies, the tissue-coverslips were post-fixed with 1.6% PFA for 10 min at RT, washed, post-fixed with ice-cold methanal for 5 min and washed. The tissue-coverslips then could be stored in the storage buffer for 5 days at 4°C or immediately loaded into CODEX instrument for multiple cycle immunostaining. The CODEX fluidics and Keyence microscope was set up using CODEX instrument manager (CIM) software and Keyence software according to manufacturer’s protocol. The master mix of fluorescently-labeled-oligonucleotide probes (fluorescent-reporters) that corresponded to DNA-barcoded antibodies for multiple cycle immunostaining was prepared in a 96-well plate according to CODEX manual. Each cycle of CODEX immunostaining consisted of three steps of adding nuclear staining DAPI, Atto550-, Cy5- and Cy7-fluorescent-reporters, imaging, and removing fluorescent-reporters. Two blank cycles (DAPI nuclear stain only without any fluorescent-reporters) were ran for evaluating the level of autofluoresence and subtracting background using CODEX processor software.

### RNAscope ISH

RNAscope ISH (RNAscope) was performed according to the manufacturer’s protocol and our previously published method(28, 29) with slight modifications. All the procedures including deparaffinization, hydration and antigen retrieval were conducted to minimize RNase contamination. The tissue-coverslips were deparaffinized and hydrated as described in the CODEX section above and incubated in 3% hydrogen peroxide for 10 min at RT. After Washing in mili-water (AQUA SOLUTION, 2700PRD), the tissue-coverslips were subjected to antigen retrieval by boiling for 15 min in RNAscope^®^ Target Retrieval Reagent solution (Cat # 322000, ACD) in a 50ml glass beaker on Hotplate to undo the cross-linking. After treatment with RNAscope® Protease Plus (Cat # 322330, ACD) at 40°C for 20 min, tissue-coverslips were hybridized with RNAscope® Probe-SIVmac239 (anti-sense, Cat# 312811, ACD) at 40°C for 2 hrs. The signals were amplified and detected with RNAscope® 2.5 HD assay-Red kit (Cat# 322360, ACD), where the fast red can be visualized both in regular light and in far-red fluorescent channel. The RNAscope® negative control probe-DapB (Cat# 310043, ACD) was used as negative control.

### Combination of CODEX immunostaining with RNAscope ISH

Two different combinations, RNAscope and subsequent CODEX as well as CODEX and subsequent RNAscope, were tested. For RNAscope and subsequent CODEX approach, tissue deparaffinization, hydration, antigen retrieval, proteinase digestion, probe hybridization, signal amplification, and development were conducted by following the single RNAscope protocol as described above. After completed the RNAscope, tissue-coverslip underwent CODEX antigen retrieval as described above by following single CODEX protocol. For the CODEX and subsequent RNAscope approach, after the completion of last cycle of CODEX by following the single CODEX protocol as described above, the tissue-coverslip was removed from the CODEX instrument and continued with the single RNAscope procedure from 3% hydrogen peroxide incubation step on. After the completion of RNAscope procedure, the tissue-coverslip was stained with DAPI (1:2000) for 5 min at RT, washed, and loaded into the CODEX instrument for capturing RNAscope fluorescent image. To integrate RNAscope image to CODEX images, Keyence microscope was set using Keyence software as described above and a new blank cycle was added, where Cy5-channel was assigned to RNAscope. After capturing the RNAscope fluorescent image, image data was transferred into the CODEX file location, where raw data of CODEX plus RNAscope were processed by CODEX processor software.

## DISCUSSION

Simultaneous detection of highly multiplexed proteins and RNA *in situ* is important for understanding health and disease. The spatially resolved relationship of different cells populations, proteins and RNA in their native tissue structure bears crucial information for disease diagnosis, pathogenesis and treatment. CODEX represents a powerful new platform for highly multiplexed proteins (up to 60) *in situ* detection (6, 8, 20). RNAscope represents another power multiplex RNA detection system (up to 12) at a single-copy sensitivity and high specificity (19, 23), where the unique double Z probes and corresponding amplification probes enables concurrent signal amplification and background noise reduction. The combination of RNAscope with regular immunohistochemical or immunofluorescent staining to detect RNA and protein within the same tissue section has been reported by other (30, 31) and us (29). However, to our knowledge there is no reported study that combined CODEX and RNAscope together. To that end, we tested different combination methods and found CODEX and subsequent RNAscope approach worked reliably, while RNAscope and subsequent CODEX did not work for CODEX signal detection. The following two factors in the RNAscope procedure may damage epitopes for CODEX detection. First, RNAscope uses protease to digest the tissue to facilitate probes access to target sequences. The protease treatment has been demonstrated to be harsh for many antigenic sites, though some epitopes may survive this treatment (32). Another factor is the dehydration step in the RNAscope method. In a typical protocol, after antigen retrieval, the slide will be dip in ethanol and air dry before adding probes for hybridization. However, preservation immune-recognizable epitopes in tissues requires maintaining a certain amount of water within tissues after antigen retrieval (33, 34). Possible solution for this negative effect of dehydration can be overcome using disaccharides to protect tissue (34) or carefully avoiding tissue drying after antigen retrieval. However, the potential damage some epitopes by protease digestion in a highly multiple protein detection system make this approach not feasible. For the CODEX and subsequent RNAscope approach, we refined antigen retrieval procedure after finding it was crucial for RNAScope signal detection. Using the hotplate antigen retrieval method is easier to eliminate RNase contamination by baking glass-beaker and coverslip holder. The Comb-CODEX-RNAscope described in this paper can be further enhanced with multiplex RNAscope ISH published recently (35, 36) to maximize the detection of interested RNA signals in addition to highly multiplexed proteins.

The cross-reactivities of 24 CODEX anti-human antibodies to rhesus macaques were also evaluated and 13 were compatible with rhesus macaques. The anti-human CD3 antibody from Akoya did not work for rhesus macaques. An anti-human CD3 antibody (Clone#: SP162, Cat#: ab245731, Abcam) was conjugated with a barcode-oligonucleotide (Cat#: 5350002, Akoya), which works for rhesus macaque tissues in CODEX.

In summary, a simple and reproducible CODEX and subsequent RNAscope protocol reports here enables spatially resolved and highly multiplexed protein and RNA *in situ* detection, which will facilitate better understanding the health and disease at single cell level in tissue spatial context.

## ACKNOWLEDGEMENTS

Author’s contributions: QL and YC designed the experiments. YC conducted most CODEX, RNAscope and combination experiments. RB conducted RNAscope experiments. YC and QL prepared the manuscript and all authors provided manuscript editing. The authors would like to thank Oliver Braubach, Najiba Mammadova and Kristin Schmidt at Akoya for their CODEX technical support. The authors also would like to thank current and previous laboratory members of QL, especially Subhra Mandal and Saroj Candra Lohani for their contribution to rhesus macaque experiments and discussion.

